# MolJam: A Multidimensional Framework for Assessing Molecular Dataset Quality and Its Impact on Machine Learning

**DOI:** 10.64898/2026.08.21.746384

**Authors:** Peng Wang, Zhaoqi Shi, Xufan Gao, Ruhong Zhou

## Abstract

High-quality molecular datasets are essential for reliable machine learning in cheminformatics and bioinformatics, yet dataset quality is rarely assessed systematically and its relationship with downstream model performance remains poorly understood. Here, we present MolJam, an open-source framework for quantitative assessment of molecular dataset quality across five dimensions—structural integrity, data quality, experimental information quality, chemical space coverage, and data distribution—using 12 standardized metrics. Application of MolJam to 11 MoleculeNet and 8 ChEMBL-derived datasets revealed widespread and heterogeneous quality issues, including undefined stereochemistry in up to 70.72% of molecules, inconsistent molecular representations, and contradictory labels. We next asked whether improving these quality metrics necessarily improves machine learning performance. Refinement of the ESOL and Lipophilicity datasets increased their MolJam quality scores but produced mixed effects on predictive performance, suggesting a competing influence of reduced dataset size. Controlled ablation experiments further demonstrated that both dataset quality and data quantity contribute to model performance and, notably, that retaining molecules with incomplete stereochemical information can outperform their removal when the resulting gain in data quantity offsets the quality penalty. Thus, molecular dataset curation cannot be reduced to maximizing data cleanliness alone but requires balancing multiple dimensions of data quality against information loss. MolJam provides a standardized framework for diagnosing molecular dataset limitations, comparing benchmark quality, and quantitatively evaluating how data curation decisions influence downstream machine learning.

## 1 Introduction

In recent years, Deep Learning (DL) and Large Language Models (LLM) have shown real promises in accelerating drug discovery by streamlining a process that traditionally took years of manual research. [1–5]. Public molecular datasets such as MoleculeNet [6], the Therapeutics Data Commons [7, 8], PubChem [9] and ChEMBL-derived collections [10– 12] that are used to train these models have therefore become the foundation for model development and benchmarking. These datasets have accelerated methodological progress by providing shared evaluation grounds, enabling direct comparisons across architectures ranging from random forest to transformer [13–16].

The quality of deep learning models heavily relies on the datasets they are trained on. [17, 18] Recently, an increasing amount of scrutiny on molecular datasets has been reported, criticizing widely used datasets listed above. [19, 20] Reported defects include invalid or unparsable SMILES strings [21], inconsistent structural representations of the same compound [22, 23], undefined stereochemistry [24], contradictory labels for identical structures [25], and distributional imbalance [26, 27]. Our analysis shows that Undefined stereochemistry affects 70.72% of molecules in one of the MoleculeNet dataset. Furthermore, label contradictions affect these widely used benchmarks, effectively confusing machine learning models and researchers alike. Alleviation methods such as combining bioactivity measurements from heterogeneous experimental sources (a common practice in constructing large-scale datasets from databases such as ChEMBL) introduce substantial noise, with nearly 65% of cross-assay IC_50_ measurements differing by more than 0.3 log units as pointed out by Landrum et al.[28].

Existing tools only partially address these problems: RD-Kit [29] provides molecular parsing and canonicalization; MolVS [30] and datamol [31] offer molecular standardization; and established curation protocols outline best practices for data refinement [24, 25, 32]. Yet no widely used tool provides a unified, quantitative assessment that jointly captures structural validity, label reliability, experimental annotation, chemical-space coverage and distributional properties. The field lacks a standardized quality-assessment instrument for molecular datasets—one that produces an interpretable, multidimensional score with defect localization, rather than a binary pass/fail verdict or an *ad hoc* checklist.

Here we present MolJam, an open-source toolkit for quantitative molecular dataset quality assessment. MolJam standardizes molecular records into traceable structural representations, computes 12 quality metrics across five dimensions, aggregates them into an interpretable 0–100 quality score, and returns diagnostic reports with visual outputs for molecular datasets. Our contributions are fourfold:

- **A quantitative, multi-dimensional quality-assessment toolkit**. MolJam scores a molecular dataset across five dimensions and 12 metrics (see Table 1) on a 0–10 scale, assessing Structural Integrity, Data Quality, Experimental Information Quality, Chemical Space Coverage and Data Distribution. The final aggregated score is reported on a 0–100 scale with detailed profiles on all dimensions and metrics, allowing users to pinpoint specific dataset issues.
- **A chemistry-aware standardization protocol with user-defined grouping granularity**. Each molecule is standardized to traceable canonical and parental forms, allowing duplication and consistency to be evaluated at a granular level chosen by the user.
- **A systematic quality assessment of public benchmarks to reveal common dataset problems**. We apply MolJam to 11 MoleculeNet and 8 ChEMBL-derived datasets and quantify recurrent dataset defects: undefined stereochemistry, representation inconsistency, label contradiction, narrow chemical-space coverage, target imbalance, etc.
- **Quantifiable basis for dataset studies, with an example to elucidate on the quality–quantity dilemma**. Using a quantifiable ablation study supported by MolJam, we perform regression experiments on Lipophilicity dataset. We show that although both dataset quantity and dataset quality positively affects machine learning model (MLM) outcomes, including low-quality data without stereochemical specifications yields better results than removing these data.

**Table 1:** Quality dimensions and metrics evaluated by MolJam.

| Dimension | Metric | What it measures |
| --- | --- | --- |
| Structural Integrity | SMILES Validity | Whether each record’s molecular identifier (mostly SMILES) can be parsed into a chemically valid molecule. |
|  | Representation Consistency | Whether records of the same compound are encoded in one consistent, canonicalized and/or parent form. This metric discriminates against datasets with multiple entries that share the same parental SMILES but with different salted, solvated, or protonated forms. |
|  | Stereochemistry Completeness | Whether stereocentres and stereogenic double bonds are fully specified. |
| Label Consistency | Binary Label Consistency | Whether structurally identical records carry the same binary label (0 or 1). |
|  | Continuous Label Consistency | Whether replicate measurements of the same compound agree for continuous labels. Outliers are counted for multiple replicate measurements. |
| Experimental Information Quality | Time Label Availability | Whether records carry time annotation (e.g. assay or publication date). |
|  | Annotation Support Quality | How complete and information-rich the experimental metadata columns describe each measurement. |
|  | Type Diversity | How many distinct kinds of experimental context are present. Users are prompted to include columns such as assay type, target name, organism name, protocol, etc. |
| Chemical Space Coverage | Chemical Diversity | How diverse the molecules are across chemical space, calculated via pairwise dissimilarity and scaffold variety. |
|  | Drug-likeness | The mean drug-likeness (QED) of the molecular population. |
| Size and Distribution | Data Size | Whether the dataset is large enough to support reliable model training. |
|  | Data Balance & Distribution | Whether binary labels are balanced or the continuous labels are normally distributed. |

## 2 Results

### 2.1 MolJam Dataset Quality Assessment Protocol

MolJam operates as a five-stage pipeline (Fig. 1): data input, standardization, scoring, aggregation and penalty, and final report.

**Fig. 1:**
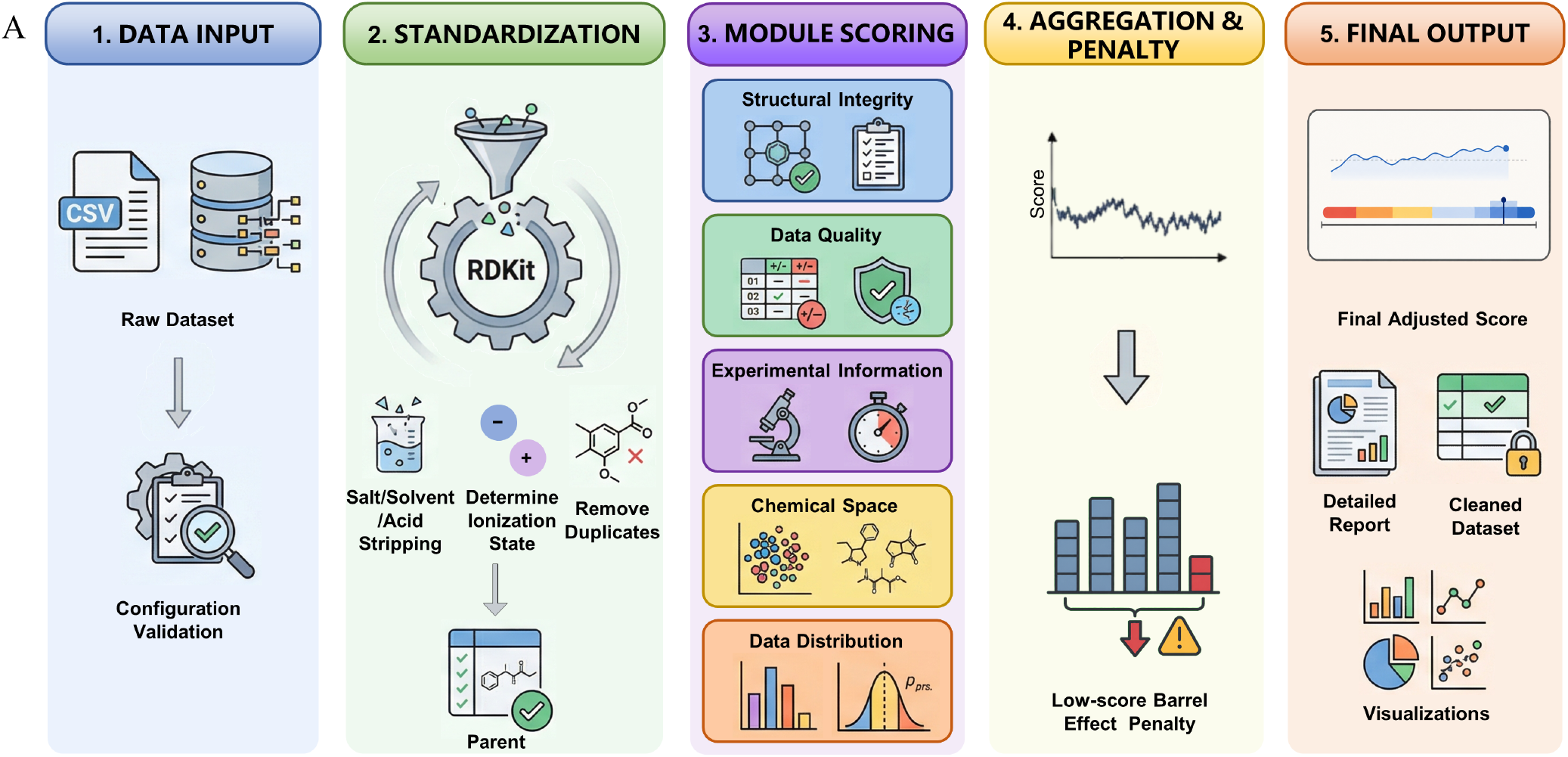
Overview of the MolJam quality-assessment framework. The pipeline operates in five stages: data input, structure standardization, module scoring,aggregation and penalty, and final output.

In data input stage, users are prompted to catogorize all columns in the tabulated dataset into three data types: molecule identifier (typically SMILES), labels, and experimental metadata. This is crucial as data columns are processed based on the assigned data types.

The standardization stage aims to detect representation consistency, label consistency and data reliability. Entries with unparsable molecule identifiers (SMILES) are labeled as invalid records. For each valid molecular record, MolJam uses RDKit to canonicalize the molecular identifier by generating `canonical_smiles`. This is followed by removing salt, solvent, and acid-adduct fragments to generate `observed_parent_smiles`. Finally, charge is removed to generate the parental SMILES `parent_smiles`. These parental forms allow MolJam to determine whether records are duplicates and should be grouped into parent-equivalent entries to be evaluated for data consistency. It is noteworthy that entries with the same parental form may refer to the same drugs with different formulation or different pH conditions. To address these concerns, MolJam provides an option to evaluate data duplication and consistency with `canonical_smiles`, `FragmentParent` or with `parent_smiles`.

The scoring stage consists of scoring functions across 5 dimensions and 12 metrics (Table 1). All metrics are mapped to a 0–10 score by one of the two functions: *Q*(*r*), a proportion-based scoring function and *C*(*c, N*), a count-based scoring function (see Methods). The exact mapping is provided in Methods.

In the aggregation and penalty stage, metrics are aggregated within each dimension and then normalized. Furthermore, a modified Cannikin Law is applied: when metrics fall below threshold values, a low-score penalty is applied to further reduce the final score. This penalty prevents a dataset with critical flaws from achieving a high overall score, even if it performs well in other metrics (Supplementary Fig. 2C–D).

Finally, MolJam returns the final score, a per-metric report, diagnostic visualizations, and optionally, a refined version of the dataset.

### 2.2 Quality Assessment of Public Molecular Datasets

We apply MolJam to 11 MoleculeNet [6] and 8 ChEMBL-derived datasets [33]. The resulting metric-level heatmap (Fig. 2A) reveals systematic patterns of quality variation across databases and dimensions.

**Fig. 2:**
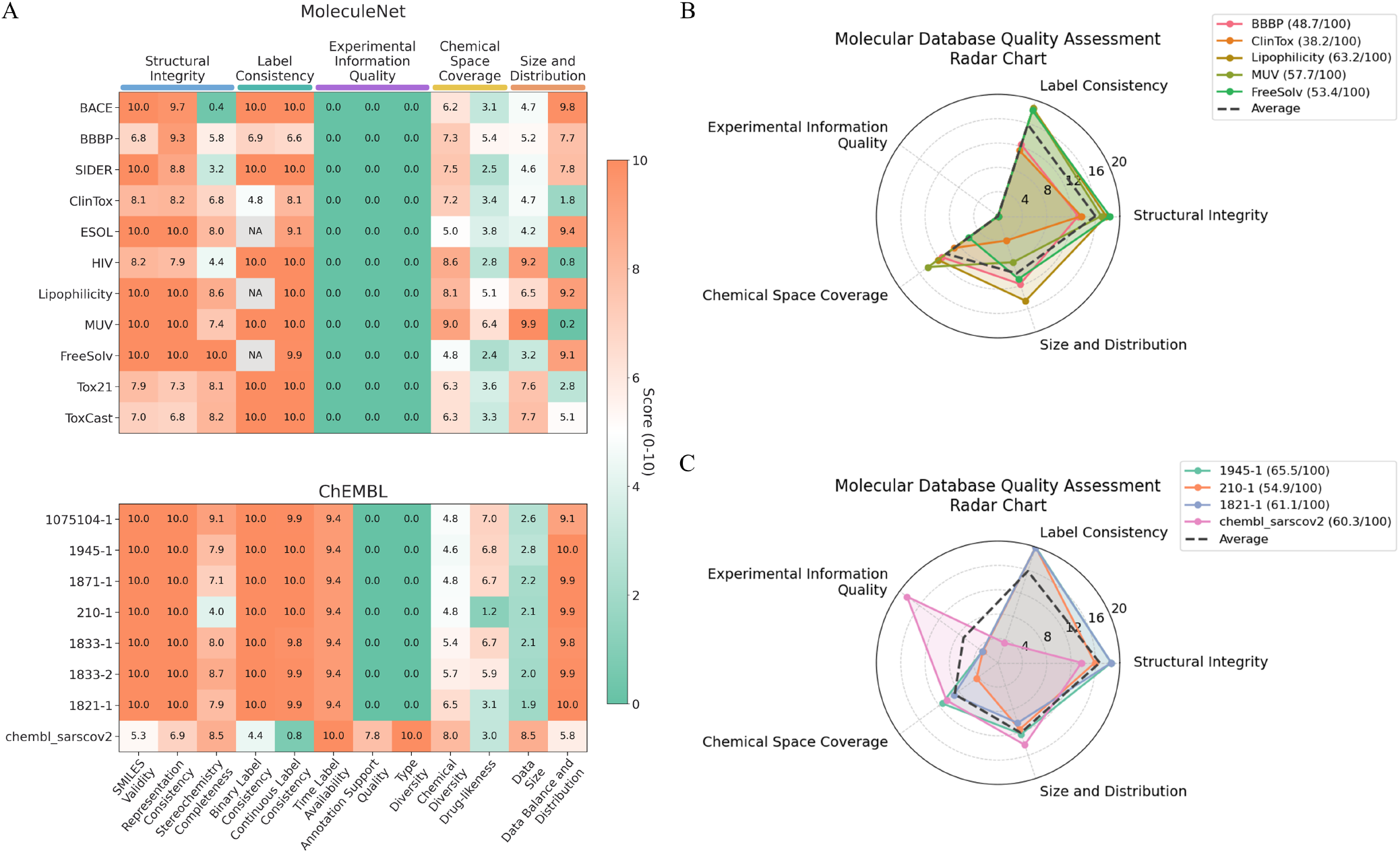
Public dataset quality profiles across 11 MoleculeNet and 8 ChEMBL-derived benchmarks. (**A**) Metric-level quality heatmap for 11 MoleculeNet and 8 ChEMBL-derived datasets. (**B**) Dimension-level radar profiles for selected MoleculeNet datasets. (**C**) Dimension-level radar profiles for selected ChEMBL-derived datasets.

#### Structural Integrity

Structral integrity issues are most prevalent in MoleculeNet datasets. SMILES Validity issues are present in five databases—ToxCast (18 entries, 0.21%), BBBP (11, 0.54%), Tox21 (8, 0.10%), HIV (7, 0.02%), and ClinTox (4, 0.27%)—with errors categorized into aromaticity, parenthesis, valence, unclosed ring, syntax, and bond anomaly types (Fig. 2B; Supplementary Fig. 3A). Stereochemistry Completeness is a dominant issue for multiple datasets: BACE exhibits the highest rate at 70.72% of molecules with undefined chiral centers (1,070/1,513), followed by SIDER (46.74%), HIV (40.81%), and BBBP (33.69%) (Table 1). Stereogenic double bonds are also widely underspecified, with HIV (23.34%) and SIDER (20.25%) most affected. For Representation Consistency, we choose `parent_smiles` for the analysis throughout this study for simplicity. It is most severe in ToxCast: 348 (4.06%) groups of molecules that share the same `parent_smiles` have inconsistent representations, 59.6% of which are salt-related duplicate entries. In Tox21, 209 (2.67%) groups of molecules that share the same `parent_smiles` have inconsistent representations, 43.1% of which are duplicates with variant protonation states (Fig. 2C; Table 2).

**Table 2:** Quality issues across MoleculeNet datasets. Numbers in parentheses indicate percentages.

| Dataset | *N* | Invalid SMILES | Undefined Chiral Centers | Stereogenic Double Bond | Structural Duplication | Label Contradiction |
| --- | --- | --- | --- | --- | --- | --- |
| BACE | 1513 | 0 | 1070 (70.72%) | 36 (2.38%) | 0 | 0 |
| BBBP | 2050 | 11 (0.54%) | 687 (33.69%) | 406 (19.91%) | 64 (3.14%) | 10 (0.51%) |
| SIDER | 1427 | 0 | 667 (46.74%) | 289 (20.25%) | 0 | 0 |
| ClinTox | 1484 | 4 (0.27%) | 402 (27.16%) | 231 (15.61%) | 19 (1.28%) | 19 (1.30%) |
| ESOL | 1128 | 0 | 212 (18.79%) | 111 (9.84%) | 11 (0.98%) | NA |
| HIV | 41127 | 7 (0.02%) | 16782 (40.81%) | 9598 (23.34%) | 0 | 0 |
| Lipophilicity | 4200 | 0 | 592 (14.10%) | 97 (2.31%) | 0 | NA |
| MUV | 93087 | 0 | 21437 (23.03%) | 6117 (6.57%) | 0 | 0 |
| FreeSolv | 642 | 0 | 1 (0.16%) | 22 (3.43%) | 0 | NA |
| Tox21 | 7831 | 8 (0.10%) | 1412 (18.05%) | 789 (10.09%) | 0 | 0 |
| ToxCast | 8597 | 18 (0.21%) | 1484 (17.30%) | 849 (9.90%) | 0 | 0 |

**Table 3:**
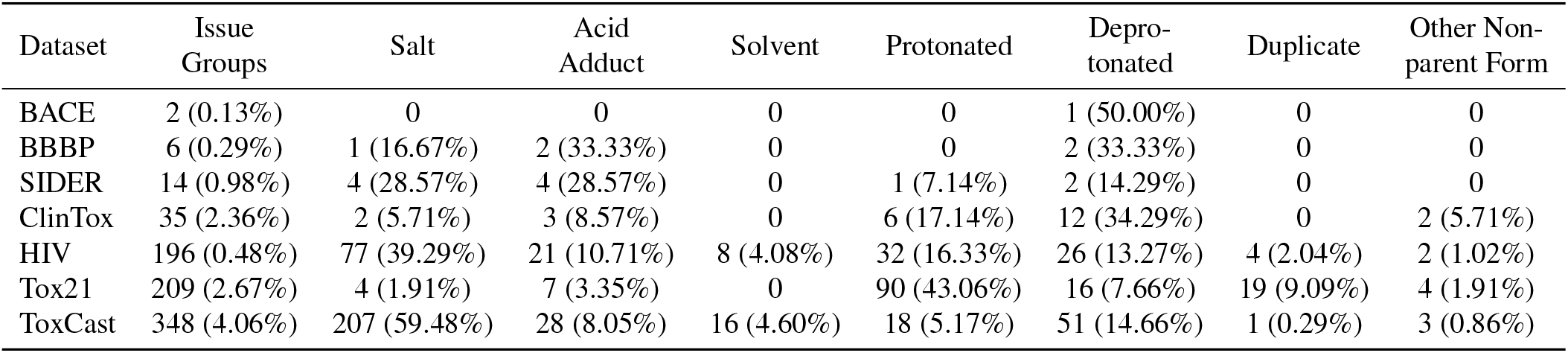
Representation consistency issue breakdown. Percentages are relative to the number of issue groups within each dataset. Datasets with zero issues are omitted.

#### Label Consistency

Note that Label Consistency is separated from Representation Consistency to capture distinguishable issues: Representation Consistency detects whether a dataset has standardized all entries, while Label Consistency detects whether a dataset has resolved data inconsistency or data contradictions. These issues are concentrated in specific datasets. Binary label consistency, measuring identical canonical structures mapped to conflicting binary labels, affecting ClinTox (19 groups, 1.30%) and BBBP (10 groups, 0.51%) (Fig. 2E). Continuous data consistency, measured by detecting outliers among replicate measurements, affects ESOL (Fig. 2F).

#### Experimental Information Quality

This dimension sharply differentiates the two dataset families: all 11 MoleculeNet datasets lack experimental-metadata columns and therefore score zero across the three metrics. In contrast, ChEMBL-derived datasets are positively scored in this dimension because of the availability of experimental meta-data. More specifically, `simpd` ChEMBL-derived datasets [33] retain time label information, and in-house dataset `chembl_sarscov2` retains time label information, experimental assay information, target organism information, among others commonly found in ChEMBL [12]. Because of this, all ChEMBL-derived datasets are given high scores for having time labels. Annotation Support Quality and Type Diversity are scored based on information quality and richness. Both scores remain 0 for the `simpd` datasets because all experimental metadata other than time labels were excluded during data curation [33]. Meanwhile, the in-house `chembl_sarscov2` dataset is properly scored due to the retainment of all experimental metadata columns from ChEMBL.

#### Chemical Space Coverage

Chemical space coverage varies dramatically across datasets. Chemical Diversity scores poorly for ESOL and FreeSolv. This is evidenced by the narrow distribution of these two datasets in t-SNE projections of Morgan fingerprints for all MoleculeNet datasets (Fig. 3G; Supplementary Fig. 4) [34, 35], which is consistent with their low scaffold diversity—the top scaffold alone accounts for 28.10% of ESOL and 49.84% of FreeSolv [36] (Fig. 3F). In contrast, MUV distributes broadly in t-SNE plot with its largest scaffold only occupying 2.24% of the dataset. Drug-likeness [37], measured by the mean QED score, also penalizes the scores of FreeSolv and ESOL (Fig. 2H), whose molecular populations include a high proportion of compounds with low drug-like properties.

**Fig. 3:**
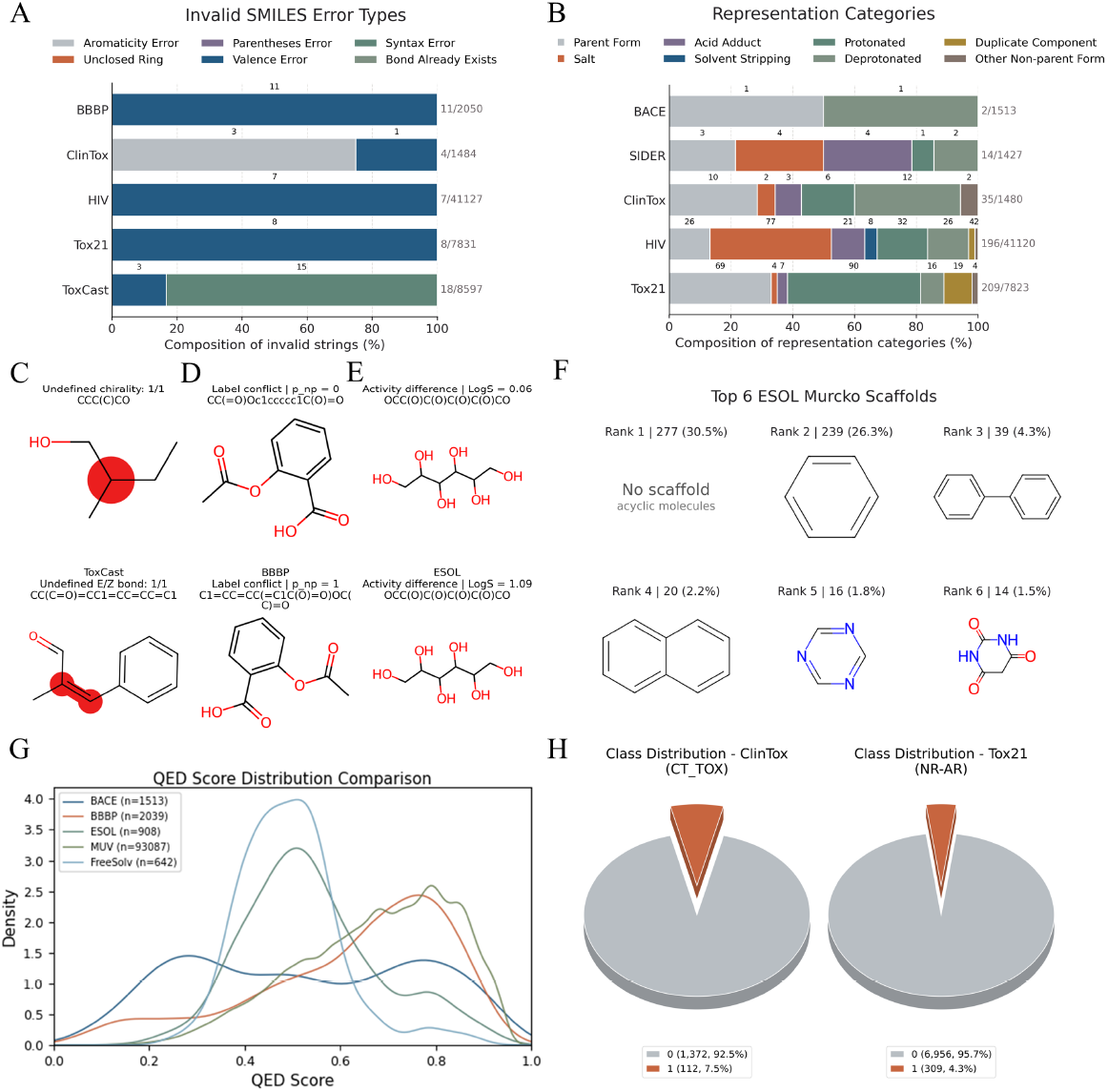
Representative structural issues, label consistency issues and distributional issues from public datasets. (**A**) Invalid SMILES error-type composition. (**B**) Representation Consistency error-category composition. (**C**) Examples of Stereochemistry Completeness issues. (**D**) A representative case of binary label inconsistency. (**E**) A representative case of continuous label inconsistency. (**F**) Top ESOL Murcko scaffolds. (**G**) QED score distributions. (**H**) Representative cases of unbalanced Data Distributions.

#### Size and Distribution

With the simple logarithm-based approach, small datasets such as FreeSolv (642 molecules) and ESOL (1,128) score poorly and large datasets such as HIV (41,127) and MUV (93,087) score more favorably. As for Data balance, datasets with even distribution are more favored by the scoring function for binary labels, and datasets with normal distribution are favored by the scoring function for continuous data. For example, binary dataset BACE with nearly equal distribution of positives and negatives is scored favorably (positive label 45.7%). Meanwhile, significantly imbalanced datasets such as HIV (3.5% active), ClinTox (7.5% toxic) are scored unfavorably. The multi-label datasets Tox21 and MUV panels are the most imbalanced, where scores are calculated for all labels and averaged. In contrast, the continuous datasets are generally scored favorably due to their near–normal distributions: ESOL, FreeSolv, and Lipophilicity (where mean–median skews are within *±*0.15 standard deviations). Specifically, the Lipophilicity dataset spans *−*1.5 to 4.5 log units with broad coverage and limited mean–median skew, resulting in a favorable Data Balance and Distribution evaluation.

Overall, Fig. 2A shows the heatmap provided by MolJam which reveals distinct shortcomings and strong suits for different datasets.

### 2.3 Aggregated MolJam scores and Multi-dimensional Quality Profiles

Datasets can be evaluated at the level of dimensions and the overall aggregated MolJam score. To compare datasets at the level of dimensions rather than individual metrics, we condense the five dimension scores into radar profiles for representative MoleculeNet (Fig. 2B) and ChEMBL-derived (Fig. 2C) datasets. All scores are normalized so that each dimension scores up to 20 points. Furthermore, a modified Cannikin Law is applied to scores for each metric to emphasize the shortcomings of a dataset (see Methods – Aggregation and low-score penalty for details). For example, Lipophilicity dataset receives deductions from Time Label Availability, Annotation Support Quality, Type Diversity and Drug Likeness, resulting in a 4.7 point penalty. Finally, scores from all dimensions are aggregated with low-score penalty applied to yield the aggregated MolJam scores (Fig. 2B and Fig. 2C).

The aggregated scores are provided as guidance while the radar profiles are provided to diagnose shortcomings of datasets. For example, Lipophilicity dataset and MUV dataset score similarly (63.2/100 and 57.7/100, respectively) despite their different radar profile. Lipophilicity, the highest-scoring MoleculeNet dataset at 63.2/100, owes its lead to high Structural Integrity and Size and Distribution. Meanwhile, MUV (57.7/100) scores the highest in Chemical Space Coverage of any dataset but scores poorly on Size and Distribution. At the lower end, BBBP (48.7/100) scores poorly in Label Consistency and Structural Integrity. ClinTox (38.2/100) scores poorly in all dimensions.

The ChEMBL-derived datasets (Fig. 2C) are generally scored higher by MolJam scoring functions (>60/100 except ChEMBL 210-1). Upon inspections, they typically score highly on Experimental Information Quality, Structural Integrity, and Size and Distribution. Interestingly, because ChEMBL datasets are typically curated from a selected number of studies, each focusing on a lead series, most ChEMBL datasets score low on Chemical Space Coverage. In contrast, `chembl_sarscov2` (60.3/100) shows the a drastically different shape with the highest scored Chemical Space Coverage and Experimental Information Quality of all datasets but poorly scored Structural Integrity and Label Consistency, likely due to the high number and wide variety of assays conducted in the past 6 years in the field.

In summary, the aggregated scores quantify the overall quality of datasets, while the radar profiles and heatmaps pinpoint where the dataset may need improvement from data curation.

### 2.4 MolJam Dataset Refinement Workflow

A refinement workflow for low-quality datasets is experimented in this study, summarized in Supplementary Fig. 4. The steps are as follows: first, records with invalid SMILES are excluded. Second, records with unresolved chiral centers or stereogenic double bonds are excluded. This step is optional if the user decides to perceive these entries as mixtures of chiral or geometric isomers. Third, for binary datasets, `parent_smiles` groups with conflicting labels are removed, and records with the same `parent_smiles` and same labels are deduplicated. Fourth, for regression datasets, entries with the same `parent_smiles` and same activity labels are removed.

We test the refinement workflow on four MoleculeNet datasets. As shown in Supplementary Fig. 6, refinement of BACE and HIV increases the MolJam score by 5.9 and 5.1 points, respectively (BACE: 50.3 *→* 56.3; HIV: 48.3 *→* 53.3; Supplementary Fig. 6), mainly through gains in Structural Integrity. Refinement of ESOL and Lipophilicity improves the MolJam scores by 2.9 points and 1.8 points, respectively (ESOL: 53.8 *→* 56.7; Lipophilicity: 63.2 *→* 65.0; Supplementary Fig. 6). More specifically, scores in SMILES Validity, Representation Consistency and Stereochemistry Completeness are all increased, as expected from the refinement procedure.

A natural question then is whether these refined datasets with higher MolJam score also lead to better machine learning model (MLM) performances. To test this hypothesis, a systematic comparison comprising 18 machine learning algorithms and 38 molecular representations is performed and discussed below.

### 2.5 Downstream MLM Performance

As shown in Fig. 4, MLMs are trained with a total of 504 combinations between algorithms and molecular representations, listed in the subplots. [5, 34, 38–40]. ESOL and Lipophilicity datasets are selected, which represents a relatively poorly scored MoleculeNet dataset (53.8 before, and 56.7 after refinement) and a relatively favorably scored MoleculeNet dataset (63.2 before, and 65.0 after refinement). Because both are continuous datasets, regression tasks are performed and model performances are measured by RMSE.

**Fig. 4:**
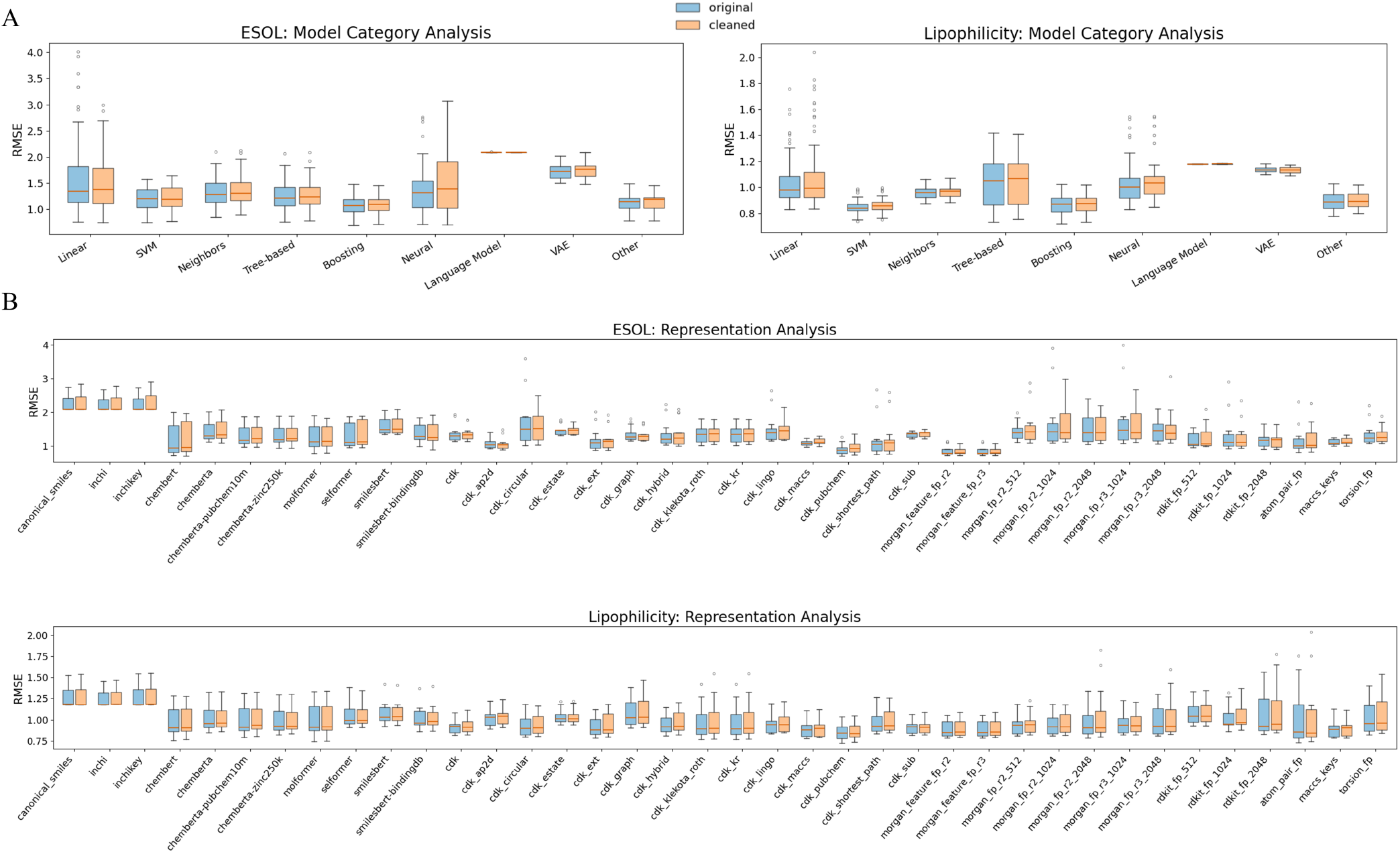
Impact of dataset refinement on downstream MLM performance. (**A**) RMSE distributions grouped by model category for ESOL and Lipophilicity before (blue) and after (orange) refinement. (**B**) RMSE distributions grouped by molecular representation for ESOL and Lipophilicity before (blue) and after (orange) refinement.

We summarize the outcome by *improvement rate*, defined as the fraction of algorithm-representation combinations with which using the refined dataset yields a lower RMSE than using the original dataset. To assess whether nominal RMSE changes are robust across random splits, paired t tests are further performed across the five random seeds for each algorithm-representation combination.

For Lipophilicity dataset, the improvement rate is 21.4%: refinement lowered RMSE in 108 of the 504 algorithm-representation combinations. The median RMSE is slightly increased from 0.943 to 0.950 log units and the mean increased from 0.981 to 0.998 log units (with relative increases of 0.7% and 1.7%, respectively; Fig. 4). Consistently, RMSE distributions before and after refinement largely overlap when grouped by model category or molecular representation. Paired t tests show that most settings have no significant RMSE difference between the original and refined datasets: 353 of the 504 combinations are insignificant at *p <* 0.05, and only 8 combinations show significantly lower RMSE after refinement.

For ESOL dataset, the improvement rate is 35.7%: refinement lowered RMSE in 180 of the 504 algorithm-representation combinations. The median RMSE is changed from 1.213 to 1.229 log units and the mean from 1.330 to 1.340 log units (with relative increases of 1.3% and 0.8%, respectively; Fig. 4). Similar to Lipophilicity, the boxplots grouped by model category and molecular representation show no clear global shift after refinement. Paired t tests again indicate that significant improvement is not widespread: 378 of the 504 combinations are insignificant at *p <* 0.05, and 44 combinations show significantly lower RMSE after refinement.

Although the refinement procedures used in this study have improved dataset quality for ESOL and Lipophilicity based on MolJam scores, data sizes have decreased. Number of entries in ESOL decreases from 1,128 to 908, which corresponds to a 19.5% reduction; Number of entries in Lipophilicity decreases from 4,200 to 3,608, which corresponds to a 14.1% of the original dataset, respectively. If we assume that both dataset quality and data size positively affect downstream MLM performance, mixed results showcased here may be explained by the contradiction of the dataset quality improvement and the data size reduction. This motivates a controlled ablation study separating the impact of dataset quality and data size, as explained below.

### 2.6 Quality versus quantity – is there a clear answer to which is more important?

The experimented refinement process in this study heavily relies on reduction of low quality data to achieve overall higher MolJam scores. Such deletion can reduce data size and can also alter chemical space coverage. In some cases such as SMILES validity, it is difficult to recover the valid SMILES and therefore these entries have to be deleted. However in other cases such as Stereochemistry Completeness, researchers may have an option to assume entries to be racemic, or one of the stereochemical isomers, thus keeping the problematic entries. As a result, a common dilemma is faced by researchers when dealing with a dataset with mixed quality: is quantity more important, or is quality more important?

To tentatively probe this question, we perform a controlled ablation study starting from the refined Lipophilicity dataset (Fig. 5). For all molecules that originally carry stereochemical specification, we select an increasing fraction (0%, 5%, 10%, 15%, and 20%) of molecules under three strategies. In the *strip* strategy, stereochemical markers are stripped from the selected molecules but the molecules are retained, yielding lower-quality datasets with unchanged size (Fig. 5A). In the *remove* strategy, the same molecules are removed by the refinement rules, yielding higher-quality but smaller datasets (Fig. 5B). In the *random remove* strategy, same amount of random molecules are removed, yielding smaller datasets with largely unchanged quality scores (Fig. 5C). All experiments are repeated 3 times with different seeds to randomly select molecules to strip or remove.

**Fig. 5:**
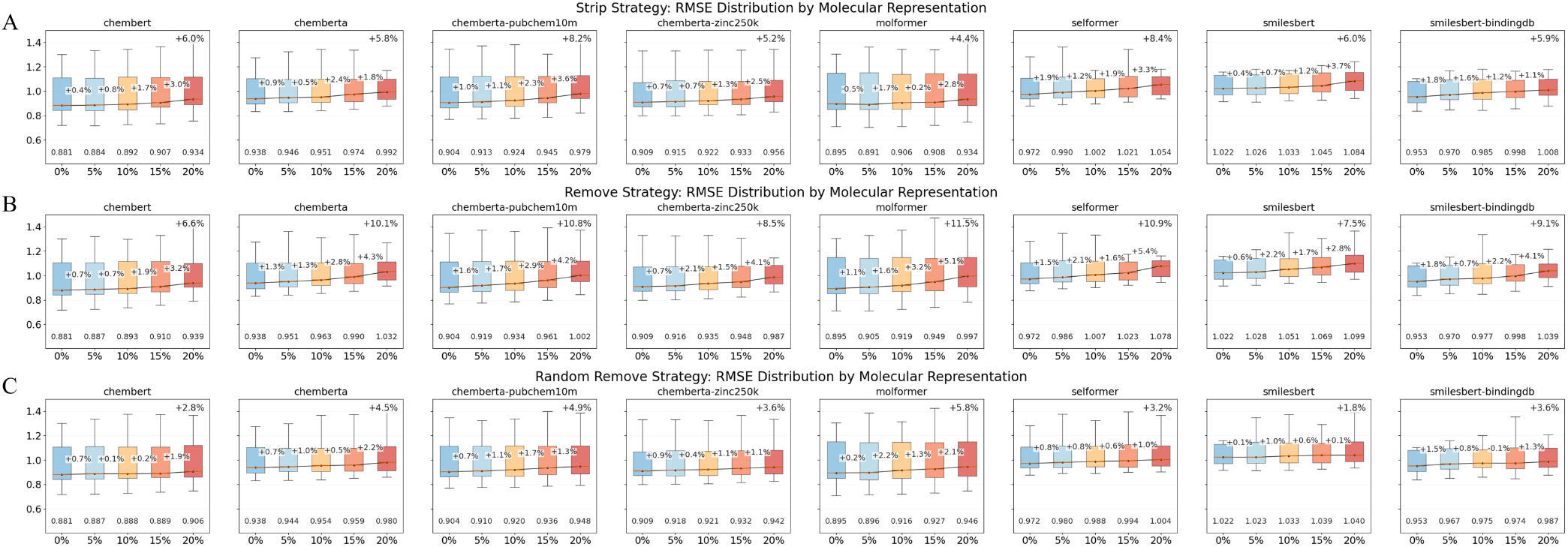
Ablation experiments controlling dataset quality and data size for Lipophilicity dataset. Starting from the refined Lipophilicity dataset, three dataset perturbation strategies are applied: (**A**) *strip* strategy: stereochemical markers are stripped from the selected molecules but the molecules are retained, yielding lower-quality datasets with unchanged size. (**B**) *remove* strategy: the same molecules are removed, yielding higher-quality but smaller datasets. (**C**) *random remove* strategy: same amount of random molecules are removed, yielding smaller datasets with largely unchanged quality scores. All experiments are repeated 3 times with different seeds to randomly select molecules to strip or remove. Boxplots show RMSE distributions across model settings, and percentage annotations indicate performance degradation relative to the starting refined dataset.

As the ablation fraction increase, performance degraded under all three strategies, indicating that both quality and quantity positively affects model performances. Specifically, the *strip* strategy (Fig. 5A) only lowers dataset quality and degrades model performance by 6.2 *±* 1.4% on average. The *random remove* strategy only lowers dataset quantity and degrades model performance by 3.8 *±* 1.3% on average. The *remove* strategy (Fig. 5B) lowers both dataset quality and dataset quantity and degrades model performance by 9.4 *±* 1.8% on average. Comparing the *strip* strategy and *remove* strategy is interesting: the degradation is consistently steeper when the stereochemical molecules are removed (Fig. 5B) than when they are stripped of stereochemical markers (Fig. 5A). This pattern seems to indicate that retaining imperfect but information-bearing records can benefit learning more than simply removing the entries, specifically for the case of Stereochemistry Completeness. It is unclear if this conclusion is generalizable as more experiments are needed for more types of datasets. Nonetheless, with MolJam quality assessment tool, we show an example to quantitatively compare the effects of quality versus quantity regarding quality dimensions such as Stereochemistry Commpleteness.

## 3 Discussion

Public molecular benchmarks are widely used to train predictive models and compare algorithms. Benchmarks are created by data mining through literature, hit/lead finding campaigns from pharmaceutical industry, or more recently, cross-disciplinary efforts designed for data curation [41–43]. Because data curation and data consumption are frequently managed by different research entities, a misalignment arises regarding the perception of dataset quality. Therefore, it is common practice to preprocess datasets to remove unparsable entries and rebalance label distribution. At the data curation stage, previous work has established the importance of data quality in cheminformatics. Fourches et al. introduced the “trust, but verify” principle for QSAR datasets [25, 32], and the ChEMBL team developed an open-source curation pipeline [24]. At the data preprocessing stage, tools have been proposed, including RDKit for parsing and canonicalization [29], MolVS for standardization [30], datamol for convenient standardization interfaces [31], and published protocols for data cleaning [24, 25]). MolJam is proposed to bridge this gap by unifying the perception of dataset quality in a consistent and interpretable manner. To the authors’ knowledge, MolJam is the first quantitative dataset evaluation tool. The aggregated MolJam score aims to provide guidance on the overall quality of the datasets. For example, MoleculeNet datasets are typically scored lower than ChEMBL datasets. This result aligns with anecdotal evidence reported in prior studies [44, 45]. In practice, MolJam can be used in combination of dataset curation protocols and dataset refinement tools to guide researchers to create datasets with higher quality, or to refine datasets for downstream ML tasks.

By evaluating datasets across 5 dimensions and 12 metrics, MolJam precisely pinpoints underlying issues. In Fig. 2, MolJam reveals the widely known issue of unparsable entries from 5 out of 10 MoleculeNet datasets. Moreover, severe problems with stereochemistry compeleteness, drug likeness, data size and data balance issues are detected across the board for MoleculeNet datasets, agreeing with the anecdotal findings reported previously [44]. Moreover, without providing any experimental metadata such as time label, data source, organism type, assay type, etc., MoleculeNet datasets are considered “incomplete” by our standards and therefore scored zeros in the dimension of Experimental Information Quality. In comparison, pre-refined ChEMBL dataset `simpd` by Landrum et al. [33] removed unparsable SMILES and data inconsistency, which is detected and confirmed by MolJam (Fig. 2A). However, such refinement protocol inevitably reduces data size and chemical diversity, which is also detected by MolJam. `ChEMBL_sarscov2` is a dataset curated in-house as a control experiment. It shows that original ChEMBL database indeed contains invalid SMILES and incosistent labels. Moreover, as pointed out above, MolJam encourages inclusion of experimental metadata in addition to time labels, as shown by the higher scores on Annotation Support Quality and Type Diversity from `chembl_sarscov2`dataset. Additional experimental metadata allows options to split data across different targets, phenotypes, organisms, assays, time and even geographic locations.

Stereochemical information is essential for property predictions, especially regarding amino acids, carbohydrates and activity cliff [46]. MolJam’s stereochemistry metric penalizes undefined stereocenters by design, aiming to encourage data curators to collect data on pure enantiomers. However, due to technological and budgetary restrictions, such data may not be obtainable. As such, entries with unlabeled stereocenters in public datasets may be considered racemates, instead of ambiguous or erroneous entries. This is a known limitation of string representations such as SMILES and SMARTS and has been a practice for data curation to some extent [47]. However, annotating stereoisomers with multiple stereocenters, especially in the cases of carbohydrates, may lead to unresolvable ambiguity in databases. For example, D-glucose and L-glucose may share the same unlabeled SMILES as `C(C1C(C(C(C(O)O1)O)O)O)O`, which contains 4 chiral centers and corresponds to 16 stereoisomers. Even for molecules with only one chiral center, presenting unlabeled data without clarification leads to unnecessary confusion. Therefore, we suggest to pair data entries with unlabeled chiral centers with an additional column of experimental metadata to specify or guesstimate the composition of the racemates.

A simple refinement protocol is developed alongside MolJam to test the impact of dataset quality on downstream ML tasks. The first sets of experiments simply remove low-quality data entries marked by MolJam and lead to mixed results. For ESOL and Lipophilicity, despite that the refined datasets yield higher final MolJam scores, downstream regression tasks do not show broadly significant RMSE improvement. These results likely indicate that higher quality scores alone do not guarantee better model performance. By refining datasets, data sizes often drop and chemical space may shift depending on the portion and type of low-quality entries removed. Therefore, in order to separate the impact of data size from data quality, we design an additional ablation study on Lipophilicity dataset, where data quality is perturbed with data size remaining unchanged. Specifically for the metric of Stereochemistry Completeness, it is shown that MLM performance deteriorates as data quality lowers. Moreover, MLM performances deteriorates more rapidly when data entries with unlabeled chiral centers are removed than when they are retained as lower-quality records. The random-deletion control shows that random removal of the same number of training molecules leads to a smaller performance loss. Firstly, these ablations studies establish that both dataset quality (by including stereochemical information) and data size positively affects MLM performances. Interestingly, these experiments also indicate that in the case of Stereochemistry Completeness, retaining stereochemically ambiguous records yields higher MLM performances by removing them, despite the best MLM performances come from the dataset with completely labeled entries. Together with our discussion above, it is important to reiterate that from the perspective of data curation, a better practice is to label racemates instead of leaving them ambiguous. However, if a data consumer for MLM is given a choice, retaining stereochemically ambiguous records may lead to better MLM performances. Such a decision must be made with cautions.

Benchmark fairness is an increasingly recognized concern in molecular ML [48–50]. Benchmark comparisons on datasets with unreported defects may favour models that tolerate noise or missing structural information rather than models that best capture structure–activity relationships. MolJam scores and metric-level reports could serve as standardized metadata accompanying published molecular datasets, enabling researchers to make informed decisions about which benchmarks are appropriate for their evaluation goals. Allowing quality profiles to accompany datasets, similar to datasheets accompanying datasets and model cards accompanying trained models [51, 52], or as the community reports model size, training cost, and energy use for model training [53, 54], introduces new standards for data curation and data governance. Such profile will clearly show whether a dataset is structurally reliable, chemically diverse, well annotated, and balanced, both for publication and consumption purposes.

Future works on MolJam may incorporate graph-level or 3D descriptors that broaden the structural metrics, and adapt the dimensional weights for aggregation. For instance, emphasizing Experimental Information Quality for datasets may benefit QSAR predictions because additional metadata may reveal hidden correlations and assist researchers to mask certain category of data. Additionally, MolJam can be paired with uncertainty-aware learning models that identify records that are most influential for model predictions, which may correlate with data entries identified as low quality by MolJam. Revealing such correlation may assist model builders to pre-filter data before resources are spent. The reliability of datasets rely much on the mutual understanding between curators and consumers. It is crucial to build infrastructures to bridge between the two communities. MolJam attempts to do so by quantifying and visualizing the quality profiles, while providing means to test more assumptions on the quality-quantity dilemma. By referring to MolJam profiles, decisions on data curation and data refinement can be made deliberately rather than anecdotally.

## 4 Methods

### 4.1 Structure Standardization

Each SMILES string is parsed with RDKit’s `Chem.MolFromSmiles` [29], and valid molecules are canonicalized with `Chem.MolToSmiles` to generate `canonical_smiles`. Thereafter, following RDKit/MolStandardize-based structure curation pipelines [24, 55], the RDKit standardization function `rdMolStandardize.Normalizer().normalize`, the Kekulization function `Chem.Kekulize`, and the canonical tautomer function `rdMolStandardize.TautomerEnumerator()` `. Canonicalize` are applied to produce `standardized_smiles`. Disconnected fragments are obtained with `Chem.GetMolFrags`, after which known solvent, salt, and acid-adduct fragments are removed using the internal SMARTS lists `_COMMON_SOLVENT_SMARTS`and `_COMMON_SALT_SMARTS` to generate `observed_parent_smiles`. For parent-level grouping, a charge-neutralized copy is generated with `rdMolStandardize.Uncharger().uncharge`, followed by the same fragment-stripping procedure to produce `parent_smiles`. Removed salts and solvents, duplicate parent fragments, parent-processing comments, fallback flags, and original indices are preserved to maintain traceability to the original entries.

### 4.2 MolJam Scoring Functions

MolJam converts raw measurement values into 0-to-10 scores using two main functions: a proportion-based scoring function *Q*(*r*) and a count-based scoring function *C*(*c, N*).

#### Proportion-based scoring function *Q*(*r*)

Given a dataset defect rate *r ∈* [0, 100%] (percentage), *Q*(*r*) returns a quality score via a three-subdomain piecewise function. Let *T*_1_ = 0.1, and *T*_2_ = 0.5 (corresponding to 10% and 50% error thresholds). Then:

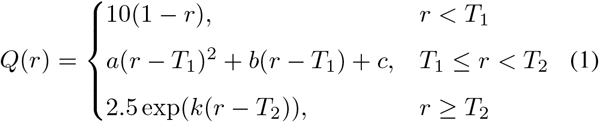

where the coefficients *a, b, c*, and *k* ensure *C*^1^ continuity between subdomains for the piecewise function (Supplementary Methods §S1). The first subdomain imposes linear scaling for low defect rates; the second subdomain utilizes quadratic polynomial equation to accelerate penalties on defect rates between 10% to 50%; and the third subdomain enforces exponential equation to further accelerate the penalties for high defect rates.

#### Count-based scoring function *C*(*c, N*)

Given a dataset defect count *c* and dataset size *N, C*(*c, N*) combines absolute and relative penalties:

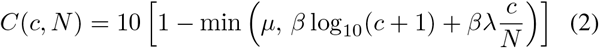

where parameters (*β, λ, µ*) are dependent on the significance of the respective metric. There are three parameter sets: the low-significance set is retained for optional count-based diagnostics but is not assigned to any of the 12 metrics in Table 1, and uses (*β, λ, µ*) = (0.05, 20, 0.5); medium-significance metrics, including Representation Consistency and the structural-duplication component of Data Consistency & Reliability, use (0.1, 50, 0.7); high-significance metrics, including Valid SMILES and Label Consistency, use (0.2, 100, 0.9) (Supplementary Table S1). Both logarithmic term on absolute count *c* and linear term on defect rate *c/N* are included in the scoring function to ensure penalty on low absolute count for severe problems (e.g., Valid SMILES and label contradictions) and linear penalty on high defect rate, with *µ* capping the total penalty.

### 4.3 Quality metrics

The 12 quality metrics listed in Table 1 use either *Q*(*r*) or *C*(*c, N*) to compute 0-to-10 scores. Below we define the input for each metric. Complete computational details are provided in Supplementary Methods.

#### Structural Integrity

##### SMILES Validity

Let *c*_invalid_ denote the count of un-parsable SMILES strings and *N* the total number of records. Then

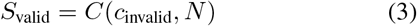

with high-significance parameters.

##### Representation Consistency

Let *c*_incon_ denote the number of inconsistent structural representations (i.e., entries with the same `X_SMILES`, where `X` can be `canonical`, `parent` etc. depending on user choice, appearing as multiple entries), and *N*_X_ the total number of `X_SMILES`. Then

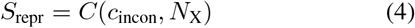

with medium-significance parameters. The details on included `X_SMILES` are provided in Supplementary Methods §S2.

##### Stereochemistry Completeness

Let *r*_chiral_ denote the percentage of molecules with at least one undefined chiral center. Then

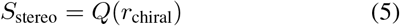

#### Label Consistency

##### Binary Label Consistency

(Binary labels only.) Let *c*_contra_ denote the number of entries with the same `X_SMILES` bearing contradictory binary labels across records, and *N*_struct_ the total number of unique `X_SMILES`. Then

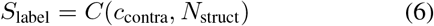

with high-significance parameters.

##### Continuous Label Consistency

(Continuous labels only.) Four sub-scores assess consistency quality among records sharing the same `X_smiles`: (i) structural duplication count *c*_dup_, scored via *C*(*c*_dup_, *N*_*valid*_) with medium-significance parameters, where *N*_*valid*_ is the number of valid entries; (ii) activity consistency, the number of duplicate entries whose coefficient of variation exceeds 0.1 (see Supplementary Methods §S3 for details); (iii) activity variability, the mean coefficient of variation 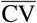 across replicate groups, scored via 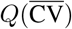; and (iv) outlier rate *f*_out_, the fraction of activity values beyond 1.5 IQR from the column quartiles, scored via *Q*(*f*_out_). All available sub-scores are then averaged to be *S*_*data*_ (Supplementary Methods §S3).

#### Experimental Information Quality

A hybrid keyword–statistical classifier identifies experimental-context columns (assay, target, biosystem, protocol, time, support metadata). For each accepted column, MolJam computes a quality signal *q ∈* [0, 1] encoding coverage, richness, and classifier confidence.

##### Time Label Availability

Let *q*_time_ denote the quality signal for time-related columns (combining coverage and richness; Supplementary Methods §S4). Then

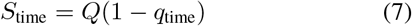

##### Annotation Support Quality

Let *q*_annot_ denote the quality signal aggregating strong experimental-context columns (weighted by classifier confidence and column entropy; Supplementary Methods §S4). Then

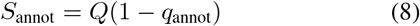

##### Type Diversity

Let *q*_type_ denote the breadth and depth across experimental-context families (Supplementary Methods §S4). Then

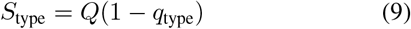

#### Chemical Space Coverage

##### Chemical Diversity

This metric sums two sub-scores (each scaled to a maximum of 5). The first is based on 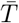, the mean pairwise Tanimoto coefficient over a fixed-seed sample of up to 1000 molecules (with Morgan fingerprints, radius 2, 2048 bits); higher similarity yields a lower score, via 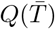. The second is based on the unique-Murcko-scaffold ratio *ρ*_scaf_ in the sample, via *Q*(1 *− ρ*_scaf_). Their sum, with a range of [0, 10], gives *S*_div_ (Supplementary Methods §S5).

##### Drug-likeness

Let 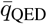 denote the mean quantitative estimate of drug-likeness (QED) [37] across valid molecules. Then

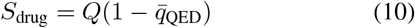

#### Size and Distribution

##### Data Size

Let *N*_*valid*_ denote the number of valid molecules. The score is computed via a seven-subdomain linear piecewise function on log_10_ *N*_*valid*_:

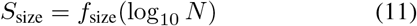

where *f*_size_ assigns 0 points for *N <* 100, linearly interpolates between breakpoints at 100, 316, 1000, 3162, 10 000, and 100 000, corresponding to *S*_*size*_ = 0, 2, 4, 6, 8, 10 and assigns 10 points for *N ≥* 100 000 (Supplementary Methods Table S2).

##### Data Balance and Distribution

For columns with continuous labels, three sub-scores assess skewness (mean-median deviation normalized by standard deviation), bin occupancy (fraction of histogram bins with non-zero counts), and range adequacy (effective span relative to expected spread). Bins are scaled based on activity units: for concentration units, bins are on a logarithmical scale; for energy units and proportions, bins are on a linear scale. All three sub-scores are computed via *Q*(*r*) and then averaged to give *S*_*bal*_. For columns with binary labels, deviation from 50:50 balance is calculated. For multiclass columns, the mean absolute deviation from uniform distribution is calculated. Both are scored via *Q*(*r*) (Supplementary Methods §S6) to give *S*_*bal*_. When both continuous and binary columns are present, their *S*_*bal*_ scores are averaged.

### 4.4 Aggregation and low-score penalty

Each dimension is normalized so that the five core dimensions (Structural Integrity, Label Consistency, Experimental Information Quality, Chemical Space Coverage, and Size and Distribution) contribute up to 20 points each; the normalized scores sum to a raw score *S*_raw_ *∈* [0, 100].

A low-score penalty then enforces a modified Cannakin Law across the 12 individual metric scores *s*_*i*_ *∈* [0, 10]. Each metric incurs a penalty

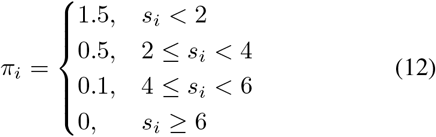

Let *ρ* denote the fraction of metrics scoring below 6. When *ρ >* 0.3, an additional penalty 1.2(*ρ −* 0.3) is added, and the total penalty is capped at 30:

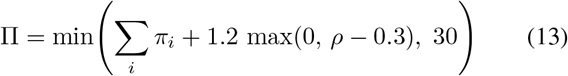

The final adjusted score subtracts this penalty from the raw score:

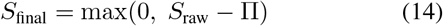

This ensures that a dataset with one or more critically weak metrics does not achieve a high final score.

### 4.5 Refinement workflow

The refinement workflow applies four sequential steps informed by the quality assessment: The steps are as follows: first, records with invalid SMILES are excluded. Second, records with unresolved chiral centers or stereogenic double bonds are excluded. This step is optional if the user decides to perceive these entries as mixtures of chiral or geometric isomers. Third, for classification datasets, `canonical_smiles`groups with conflicting labels are removed, and records with the same `canonical_smiles` and same labels are deduplicated. Fourth, for regression datasets, entries with the same `canonical_smiles` and same activity labels are removed.

### 4.6 Benchmark protocol

#### Quality assessment

We evaluate 11 MoleculeNet datasets (BACE, BBBP, SIDER, ClinTox, ESOL, HIV, Lipophilicity, MUV, FreeSolv, Tox21, ToxCast) [6], 7 ChEMBL-derived datasets [33] and 1 in-house ChEMBL dataset `chembl_sarscov2`.

#### Downstream regression

For ESOL and Lipophilicity, we compare the original and refined datasets under random train– test splits (80/20) using MLMs. 18 regression models including tree-based (decision tree, random forest, extra trees, bagging), boosting (gradient boosting, LightGBM, XGBoost), linear (ridge, Bayesian ridge), kernel (SVM-RBF, SVM-linear), neighbour (k-NN), neural-network (MLP), transformer (small, medium), and variational-autoencoder (compact, deep, latent 64/128/256) families, are combined with 38 molecular representations including ECFP, MACCS, RDKit, and Avalon fingerprints, SMILES, and SELFIES encodings [34, 38, 39, 56] for machine learning training tasks. Performances are measured by RMSE. Statistical significance is assessed by paired t tests over five random seeds for each matched model– representation configuration.

#### Lipophilicity ablation

Starting from the refined Lipophilicity dataset, we select an increasing fraction *r*_*chiral*_ (0%, 5%, 10%, 15%, and 20%) of molecules under three strategies, explained in Results. Performances are evaluated across the same 504 model–representation combinations under the random-split protocol.

## 5 Data and Code Availability

MolJam is available as an open-source Python package at https://github.com/TheZombie0/MolJam/releases/tag/v0.1.0. All MoleculeNet datasets are publicly available through the MoleculeNet benchmark [6]. ChEMBL-derived benchmark datasets are available through the `simpd` repository [33]. `ChEMBL_sarscov2` dataset is available at the same repository. The ChEMBL database is available at https://www.ebi.ac.uk/chembl/ [12]. All scoring results, refined datasets and analysis scripts used in this study are available in the same repository.

