## Supplementary Figures for "MolJam: A Multidimensional Framework for Assessing Molecular Dataset Quality and Its Impact on Machine Learning"

A

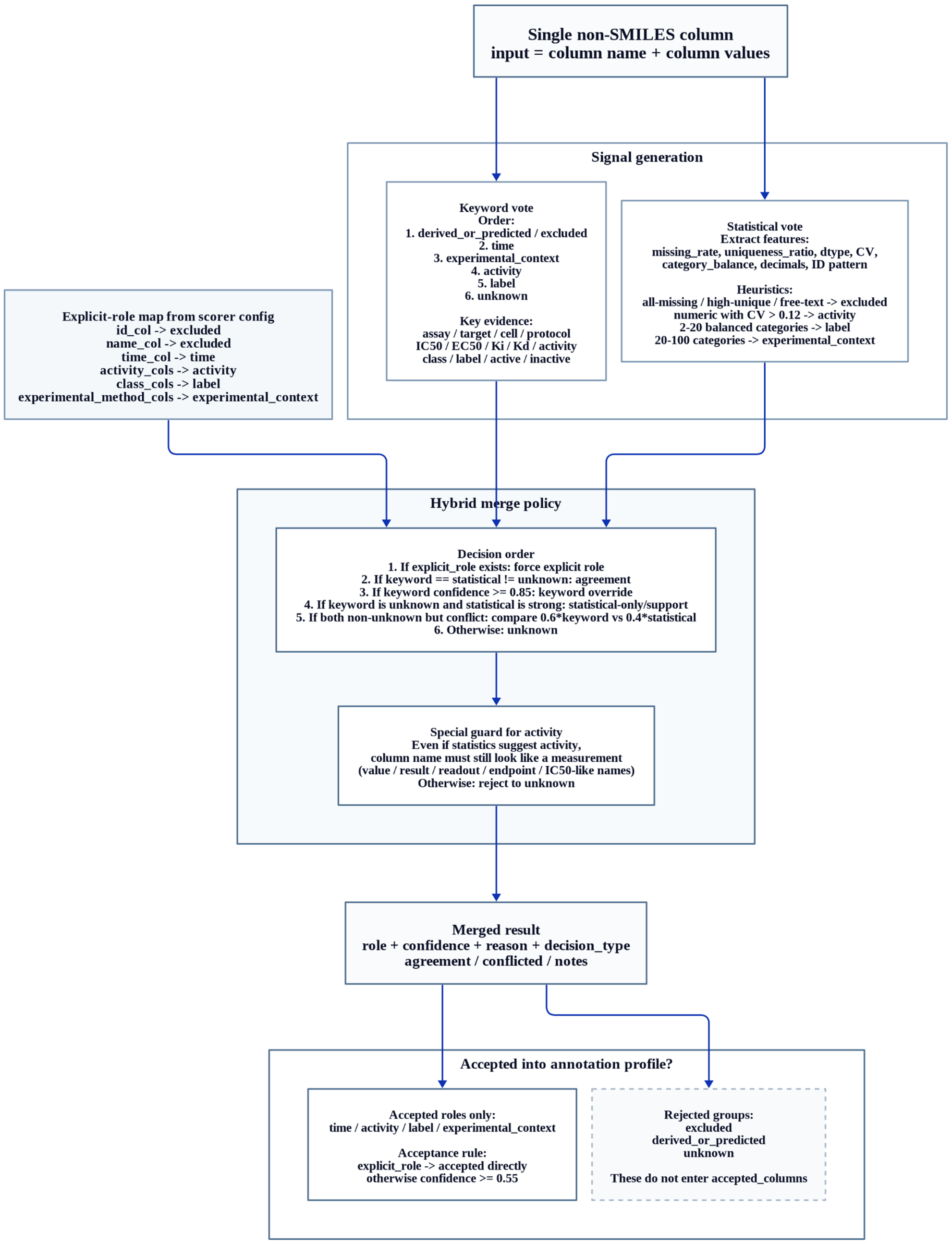

Supplementary Fig. S1: **Hybrid experimental-annotation column classifier.** Each non-SMILES column is evaluated with its column name and values. Explicit columns mappings can be supplied by users in the scorer configuration which take priority, after which a keyword vote system and a statistical vote system are applied. The keyword vote branch searches for keywords associated with time, experimental context, activity, labels, excluded columns, or unknown columns; the statistical vote branch detects heuristics such as missingness, uniqueness ratio, data type, coefficient of variation, category balance, decimal patterns, and identifier-like patterns. A hybrid merge policy combines the two signals, applies a special safe-guard rule for activity-like measurements and labels these measurements as activity and/or labels. Meanwhile, columns identified as time and experimental-context are deemed as accepted columns. Other columns are rejected and exluded from all evaluations.

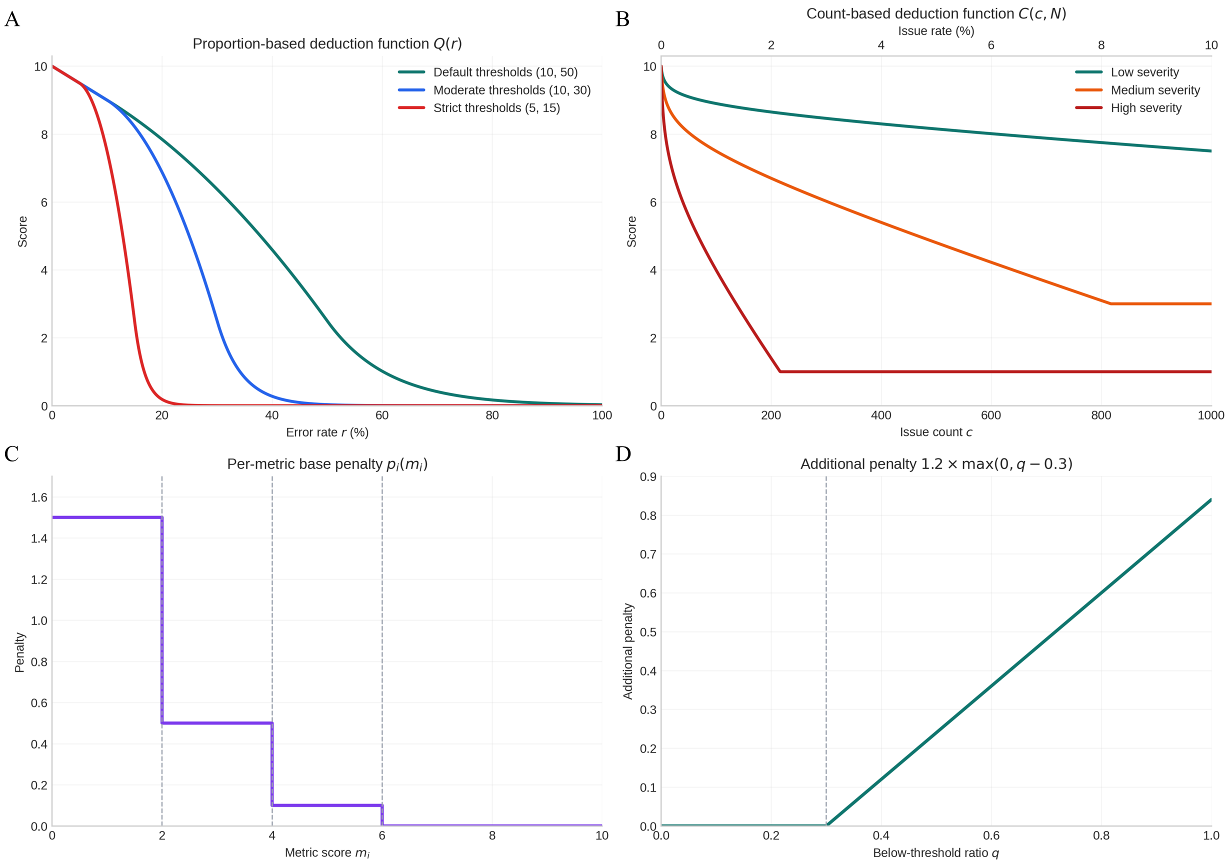

Supplementary Fig. S2: **Core scoring functions and low-score penalty mechanism.** (**A**) Proportion-based scoring function $Q(r)$ under default, moderate, and strict threshold settings. Low error rates receive a linear penalty, intermediate error rates receive an accelerated quadratic penalty, and high error rates enter an exponential decay region. (**B**) Count-based scoring function $C(c, N)$ under low-, medium-, and high-severity parameter settings, combining absolute issue count and issue rate. (**C**) Per-metric base penalty applied to individual metric scores below 6. (**D**) Additional penalty applied when more than 30% of evaluated metrics fall below the threshold.

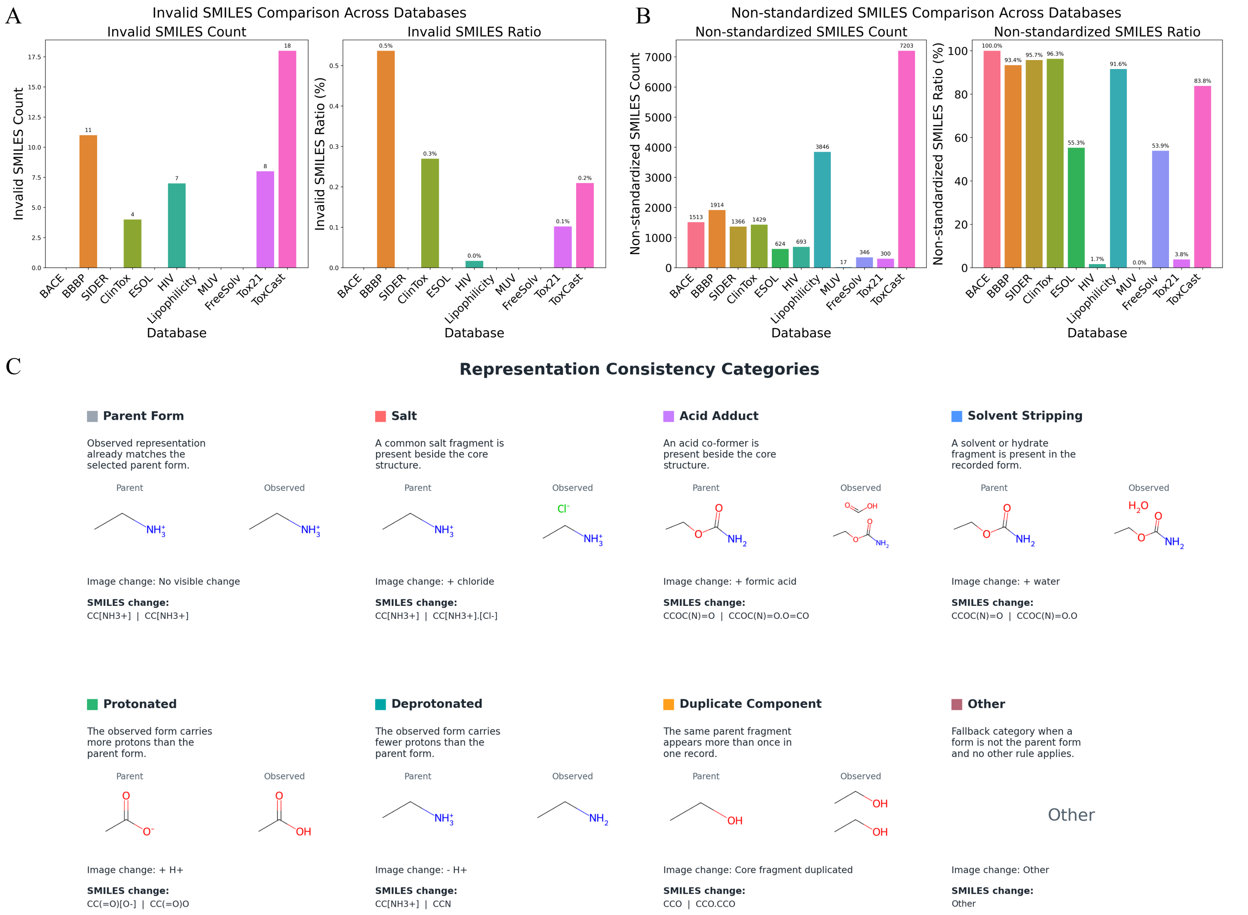

Supplementary Fig. S3: **Structural integrity diagnostics and representation categories.** (**A**) Invalid-SMILES counts and ratios across MoleculeNet datasets. Invalid SMILES are uncommon but present in ToxCast, BBBP, Tox21, HIV, and ClinTox. (**B**) Non-standardized SMILES counts and ratios, showing that raw records often differ from canonicalized representations even when they are parsable. (**C**) Representation-consistency categories used to interpret parent-level discrepancies, including parent form, salt, acid adduct, solvent stripping, protonated form, deprotonated form, duplicate component, and other non-parent form.

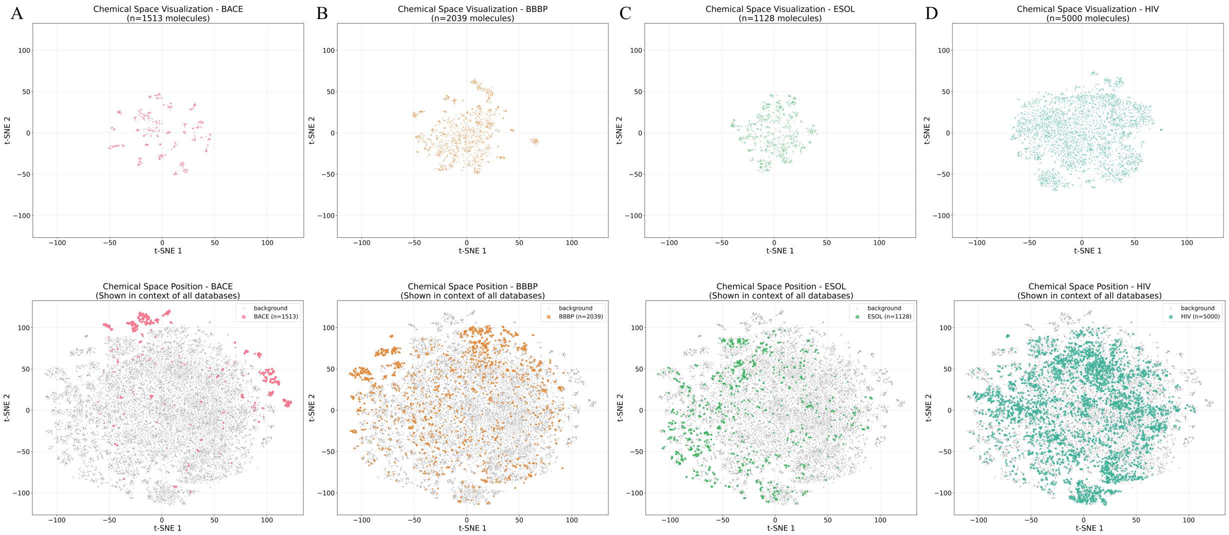

Supplementary Fig. S4: **Chemical space visualization for representative MoleculeNet datasets.** t-SNE projections of Morgan fingerprints for BACE, BBBP, ESOL, and HIV. The top row shows the distribution of each dataset alone; the bottom row places the same molecules on a combined MoleculeNet background to show their relative chemical-space positions. BACE and FreeSolv occupy narrower regions, whereas HIV spans a broader region of the combined space.

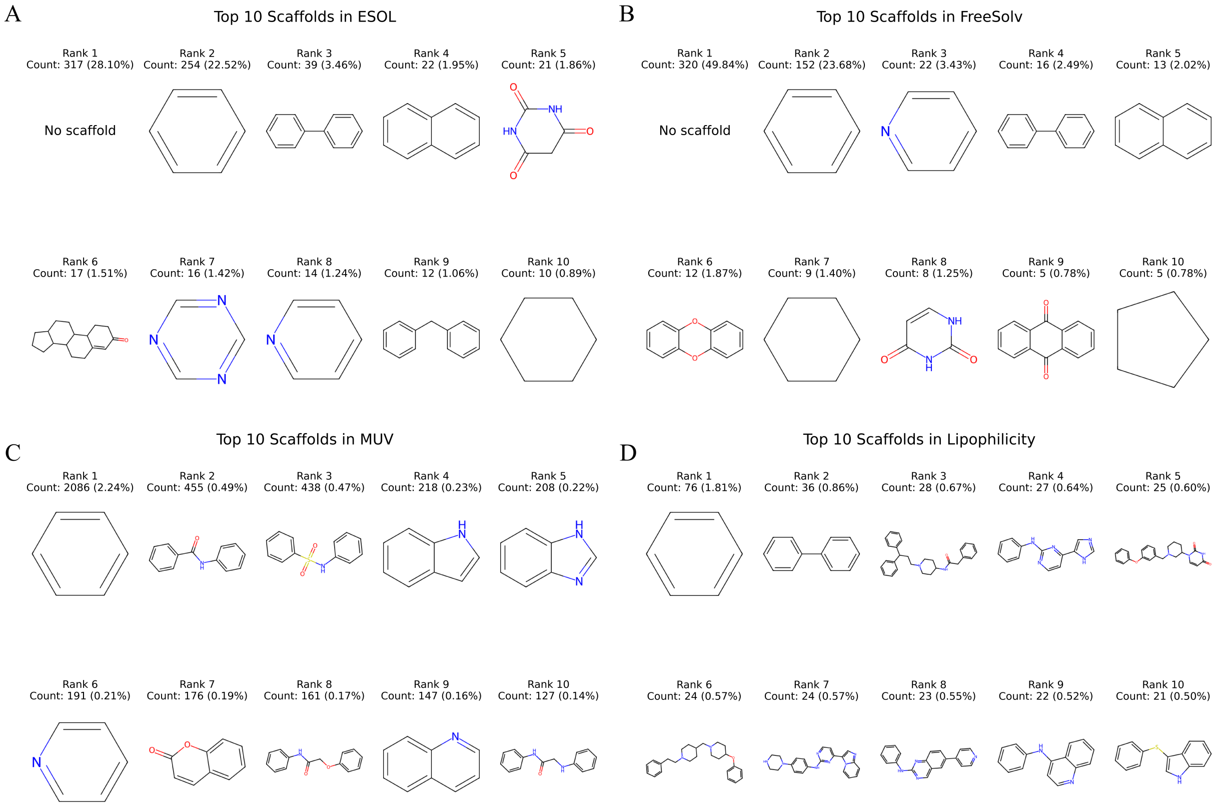

Supplementary Fig. S5: **Top Murcko scaffold profiles in selected MoleculeNet datasets.** Top 10 Murcko scaffolds are shown for ESOL, FreeSolv, MUV, and Lipophilicity. ESOL and FreeSolv are dominated by no-scaffold or simple-scaffold structures, with the largest no-scaffold group covering 28.10% and 49.84% of the datasets, respectively. MUV and Lipophilicity show more distributed scaffold profiles, with the most frequent scaffold covering only 2.24% and 1.81%, respectively.

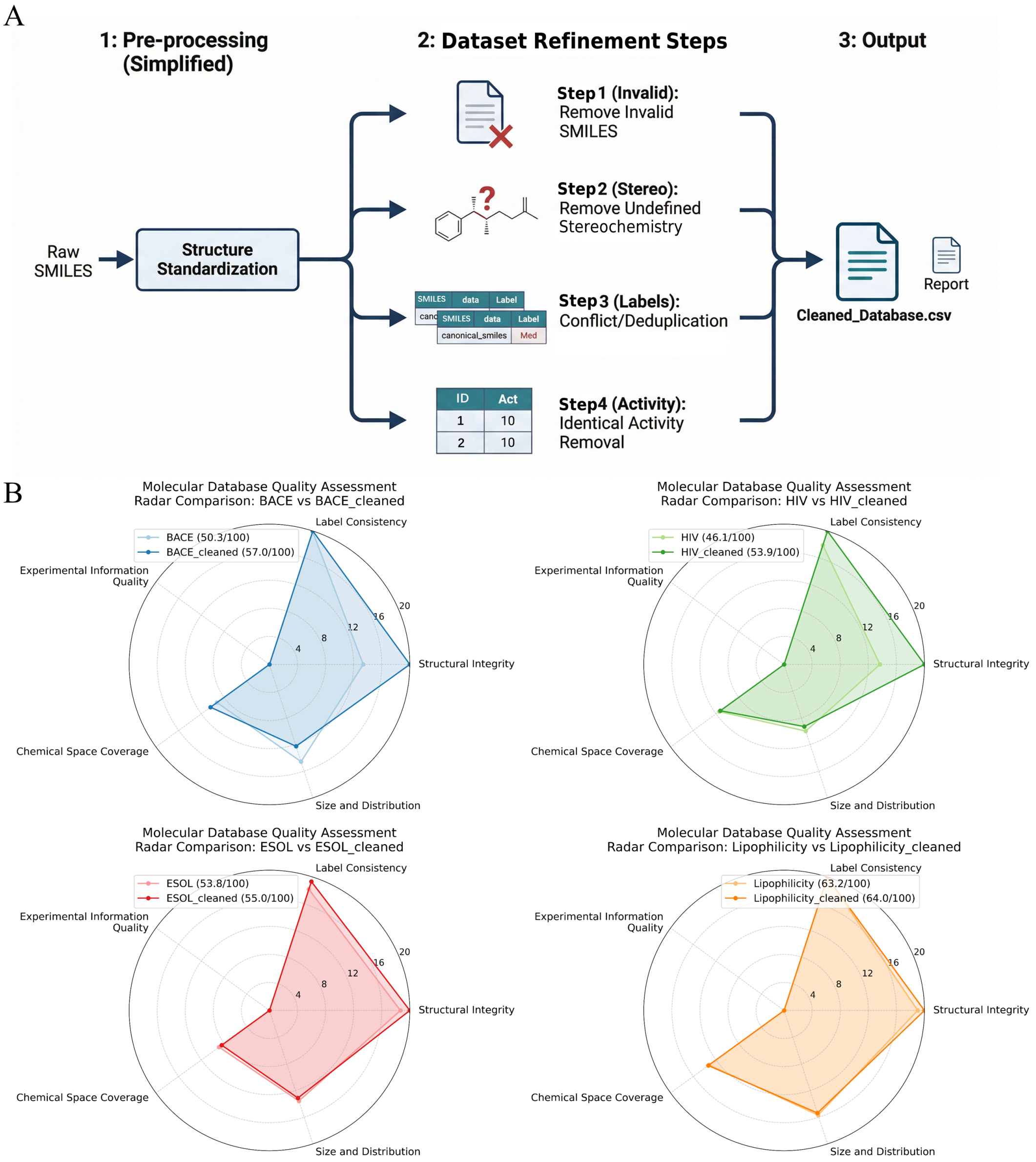

Supplementary Fig. S6: **4-step refinement workflow and quality-score changes.** (**A**) The refinement workflow first standardizes raw SMILES, then follows four steps: removal of invalid SMILES, removal of records with undefined stereochemistry, resolution of label conflicts and deduplication, and removal of identical activity records. The workflow produces a refined dataset and an associated report. (**B**) Radar comparison of BACE and HIV before and after refinement. Refinement increases the MolJam quality score from 50.3 to 57.0 for BACE, from 46.1 to 53.9 for HIV, from 53.8 to 55.0 for ESOL, and from 63.2 to 64.0 for Lipophilicity, mainly through improved structural integrity.

### S1 Core scoring functions

This section provides the original computational formulas underlying the MolJam quality metrics summarized in the main-text Methods. All formulas correspond to the reference implementation.

#### S1.1 Proportion-based scoring $Q(r)$

Given an error rate $r \in [0, 100\%]$, maximum score $S_{\max}$ (default 10), $Q(r)$ returns a quality score via a three-subdomain piecewise function. Let $T_1 = 0.1$, and $T_2 = 0.5$ (corresponding to 10% and 50% error thresholds). Then:

$$Q(r) = \begin{cases} S_{\max}(1-r), & r < T_1 \\ a(r-T_1)^2 + b(r-T_1) + c, & T_1 \le r < T_2 \\ \dfrac{S_{\max}}{4}\exp(k(r-T_2)), & r \ge T_2 \end{cases} \tag{S1}$$

with coefficients defined for $C^1$ continuity. Writing $w = T_2 - T_1$:

$$c = S_{\max}(1-T_1), \tag{S2}$$
$$b = -S_{\max}, \tag{S3}$$
$$a = \frac{S_{\max}/4 - c - b\,w}{w^2}, \tag{S4}$$
$$k = \frac{2aw + b}{S_{\max}/4}. \tag{S5}$$

The result is clamped to $[0, S_{\max}]$. Segment 1 (linear) applies to low error rates, segment 2 (quadratic) accelerates the penalty, and segment 3 (exponential decay) drives the score toward zero for severe error rates.

#### S1.2 Count-based scoring $C(c, N)$

Given a defect count $c$, dataset size $N$, maximum score $S_{\max}$ (default 10), and severity parameters $(\beta, \lambda, \mu)$, let the rate be $\rho = c/N$. The penalty combines a logarithmic absolute-count term and a linear rate term, capped at $\mu$:

$$P(c, N) = \min\Big(\mu,\ \beta \log_{10}(c+1) + \beta\lambda\,\rho\Big), \tag{S6}$$

and the score is

$$C(c, N) = S_{\max}\,\big(1 - P(c, N)\big), \tag{S7}$$

clamped to $[0, S_{\max}]$; $C = S_{\max}$ when $c = 0$. The severity parameter sets are given in Table S1.

Supplementary Table S1: Severity parameters for the count-based scoring function $C(c, N)$.

| Severity | $\beta$ (base penalty) | $\lambda$ (rate multiplier) | $\mu$ (max penalty) |
|---|---|---|---|
| Low | 0.05 | 20 | 0.5 |
| Medium | 0.10 | 50 | 0.7 |
| High | 0.20 | 100 | 0.9 |

### S2 Structural Integrity metrics

Let $N$ denote the total number of records and $V$ the set of valid molecules (those whose SMILES parse successfully).

#### S2.1 SMILES Validity

Let $n_{\text{invalid}} = N - |V|$ be the number of unparsable SMILES strings. The score is

$$S_{\text{valid}} = C\big(n_{\text{invalid}},\, N\big) \quad \text{(high severity)}. \tag{S8}$$

#### S2.2 Representation Consistency

Records are grouped by the selected parent key (default `parent_smiles`). For each record $x$, a representation signature is formed from five components:

$$\text{sig}(x) = \big(\text{can}(x),\, \text{obs}(x),\, \text{salt}(x),\, \text{solv}(x),\, \text{dup}(x)\big), \tag{S9}$$

where can$(x)$ is the canonical SMILES, obs$(x)$ the observed-parent SMILES, salt$(x)$ and solv$(x)$ the removed salt and solvent fragments, and dup$(x)$ the duplicated parent fragments. A parent group is flagged as inconsistent when its records yield more than one distinct signature. With $n_{\text{incon}}$ the number of inconsistent groups and $N_{\text{parent}}$ the total number of parent groups,

$$S_{\text{repr}} = C\big(n_{\text{incon}},\, N_{\text{parent}}\big) \quad \text{(medium severity)}. \tag{S10}$$

#### S2.3 Stereochemistry Completeness

Let $r_{\text{chiral}}$ be the percentage of valid molecules carrying at least one undefined chiral centre, i.e. $r_{\text{chiral}} = |U_\chi|/|V|$ where $U_\chi$ is the set of such molecules. The score is

$$S_{\text{stereo}} = Q\big(r_{\text{chiral}};\, S_{\max} = 10,\, T_1 = 0.1,\, T_2 = 0.5\big). \tag{S11}$$

Undefined stereogenic double bonds are reported diagnostically and removed during refinement, but the score itself is driven by undefined chiral centres at the molecule level.

### S3 Label Consistency metrics

#### S3.1 Binary Label Consistency

(Binary labels only.) Records are grouped by `parent_smiles`. A group is contradictory in a given class column if its non-missing labels are not all identical. Let $n_{\text{contra}}$ be the number of parent structures contradictory in at least one class column, and $N_{\text{struct}}$ the total number of parent-SMILES groups. The score is

$$S_{\text{label}} = C\big(n_{\text{contra}},\, N_{\text{struct}}\big) \quad \text{(high severity)}. \tag{S12}$$

#### S3.2 Continuous Label Consistency

(Continuous labels only.) Records are grouped by `parent_smiles`. Up to four sub-scores are computed and averaged over those that are applicable.

**Structural duplication.** The duplicate count is

$$n_{\text{dup}} = \sum_g \big(|g| - 1\big), \tag{S13}$$

summed over parent-SMILES groups $g$, and scored by $C(n_{\text{dup}}, |V|)$ with medium severity, where $|V|$ is the number of valid records.

**Activity consistency.** For each replicate group $g$ and activity column, the coefficient of variation is

$$\text{CV}(g) = \frac{\text{std}(g)}{|\,\text{mean}(g)|} \quad (0 \text{ if } \text{mean}(g) = 0). \tag{S14}$$

Let $m$ be the number of groups with CV$(g) > 0.1$. The sub-score is $S_{\text{cons}} = \max(0,\, 10 - m)$.

**Activity variability.** With $\overline{\text{CV}}$ the mean coefficient of variation across all replicate groups,

$$S_{\text{var}} = Q\big(\overline{\text{CV}};\ S_{\max} = 10,\, T_1 = 0.1,\, T_2 = 0.3\big). \tag{S15}$$

**Outlier rate.** Using the $1.5\times$IQR rule per activity column, with $f_{\text{out}}$ the overall fraction of outlying values,

$$S_{\text{out}} = Q\big(f_{\text{out}};\ S_{\max} = 10,\, T_1 = 0.05,\, T_2 = 0.15\big). \tag{S16}$$

The Continuous Label Consistency score is the average of all available sub-scores.

### S4 Experimental Information Quality

A hybrid keyword–statistical classifier assigns each candidate column a role and computes, per column, a coverage (non-missing fraction), a classifier confidence, and an information-strength signal. Each of the three metrics converts a quality signal $q \in [0, 1]$ to a score via $Q(1 - q)$.

#### S4.1 Time Label Availability

For the resolved time column, let $\text{cov}_{\text{time}}$ be its coverage and $n_t$ the number of unique time points. The richness component is

$$\text{rich} = \min\left(1,\ \frac{\log_2(1 + n_t)}{\log_2(1 + 4)}\right), \tag{S17}$$

and the quality signal is

$$q_{\text{time}} = 0.8\,\text{cov}_{\text{time}} + 0.2\,\text{rich}. \tag{S18}$$

#### S4.2 Annotation Support Quality

For each experimental-context column, the support signal is

$$s = \text{cov}\,\big(0.8\,\text{conf} + 0.2\,\text{info}\big), \tag{S19}$$

clamped to $[0, 1]$, where cov, conf, and info are the column coverage, classifier confidence, and information strength. Columns with $s > 0.6$ qualify. With $\bar{s}$ the mean signal over $n_q$ qualifying columns,

$$\text{strength} = \bar{s}, \tag{S20}$$

$$\text{abundance} = 1 - \exp(-n_q/4), \tag{S21}$$

$$q_{\text{annot}} = 0.6\,\text{strength} + 0.4\,\text{abundance}. \tag{S22}$$

#### S4.3 Type Diversity

Context columns are classified into four core families (assay, target, biosystem, protocol) and an optional variant family. For each family, the presence signal from its columns is

$$p_{\text{fam}} = \min\left(1,\ \sqrt{\frac{\max_{\text{col}} \text{cov}}{0.3}}\right), \tag{S23}$$

(0 if the family has no columns). Let $n_c$ be the number of core families with $p_{\text{fam}} \geq 0.4$, and $\overline{p}_{\text{core}}$ the mean presence over core families with $p_{\text{fam}} > 0$. With breadth $= n_c/4$, depth $= \overline{p}_{\text{core}}$, and a variant bonus $0.05\,p_{\text{variant}}$,

$$q_{\text{type}} = \min\big(1,\ 0.45\,\text{breadth} + 0.55\,\text{depth} + 0.05\,p_{\text{variant}}\big). \tag{S24}$$

### S5 Chemical Space Coverage

#### S5.1 Chemical Diversity

This metric sums two sub-scores, each on a 0–5 scale. From a fixed-seed (42) sample of up to 1000 valid molecules with computable Morgan fingerprints (radius 2, 2048 bits), let $\bar{T}$ be the mean pairwise Tanimoto similarity. The similarity sub-score is

$$S_{\text{sim}} = Q\big(\bar{T};\ S_{\max} = 5,\ T_1 = 0.3,\ T_2 = 0.7\big). \tag{S25}$$

Let $\rho_{\text{scaf}}$ be the fraction of unique Murcko scaffolds within the sampled molecules. The scaffold sub-score is

$$S_{\text{scaf}} = Q\big((1 - \rho_{\text{scaf}});\ S_{\max} = 5,\ T_1 = 0.2,\ T_2 = 0.6\big). \tag{S26}$$

The Chemical Diversity score is $S_{\text{div}} = \min\big(10,\ S_{\text{sim}} + S_{\text{scaf}}\big)$.

#### S5.2 Drug-likeness

Let $\overline{\text{QED}}$ be the mean quantitative estimate of drug-likeness over valid molecules with computable QED. The score is

$$S_{\text{drug}} = Q\big((1 - \overline{\text{QED}});\ S_{\max} = 10,\ T_1 = 0.2,\ T_2 = 0.5\big). \tag{S27}$$

### S6 Data Distribution metrics

#### S6.1 Data Size

Let $L = \log_{10} N$, where $N$ is the number of valid molecules. The Data Size score is the seven-segment piecewise-linear function in Table S2.

Supplementary Table S2: Data Size scoring as a function of dataset size $N$ (with $L = \log_{10} N$).

| $N$ range | $L$ range | Score $S_{\text{size}}$ | Category |
|---|---|---|---|
| $\geq$ 100,000 | $L \geq 5$ | 10 | Excellent |
| 10,000–99,999 | $4 \leq L < 5$ | $8 + 2(L - 4)$ | Very Good |
| 3,162–9,999 | $3.5 \leq L < 4$ | $6 + 4(L - 3.5)$ | Good |
| 1,000–3,161 | $3 \leq L < 3.5$ | $4 + 4(L - 3)$ | Adequate |
| 316–999 | $2.5 \leq L < 3$ | $2 + 4(L - 2.5)$ | Minimal |
| 100–315 | $2 \leq L < 2.5$ | $4(L - 2)$ | Poor |
| $<$ 100 | $L < 2$ | 0 | Insufficient |

#### S6.2 Size and Distribution

Continuous label columns and binary/categorical label columns are scored separately; when both are present, the dimension score is the mean of the two.

**Continuous labels.** For each activity column with mean $\bar{x}$, median $\tilde{x}$, and standard deviation $\sigma$, the skewness proxy is $\text{sk} = (\bar{x} - \tilde{x})/\sigma$ (0 if $\sigma = 0$). Three sub-scores are summed:

$$S_{\text{skew}} = Q\big(0.5\,|\text{sk}|;\ 3.33,\ 0.25,\ 0.75\big), \tag{S28}$$

$$S_{\text{occ}} = Q\big((1 - o);\ 3.33,\ 0.2,\ 0.6\big), \tag{S29}$$

$$S_{\text{range}} = Q\big((1 - \alpha);\ 3.34,\ 0.2,\ 0.6\big), \tag{S30}$$

where $o$ is the fraction of occupied bins in a 10-bin histogram and $\alpha \in [0, 1]$ is the range-adequacy measure (effective $q_{95} - q_5$ span relative to an expected spread, with log-scale handling for positive-valued or logarithmically scaled columns). The column score is $S_{\text{skew}} + S_{\text{occ}} + S_{\text{range}}$, and continuous columns are averaged.

**Binary/Categorical columns.** For a binary column with minority-class percentage $p_{\min}$:

$$S_{\text{bin}} = \begin{cases} 10, & 0.3 \le p_{\min} \le 0.5 \\ Q\big(3(0.3 - p_{\min});\ 10,\ 0.1,\ 0.4\big), & 0.2 \le p_{\min} < 0.3 \\ Q\big(5(0.2 - p_{\min});\ 10,\ 0.2,\ 0.6\big), & p_{\min} < 0.2 \end{cases} \tag{S31}$$

For a multi-class column with $K$ classes and class fractions $\{f_k\}$, let the mean absolute deviation from uniform be $\bar{d} = \frac{1}{K}\sum_k |f_k - 1/K|$, with maximum possible deviation $d_{\max} = 1 - 1/K$. Then

$$S_{\text{multi}} = Q\left(\frac{\bar{d}}{d_{\max}};\ 10,\ 0.1,\ 0.5\right). \tag{S32}$$

Categorical columns are averaged to give the categorical sub-score.
